# Distributional Range, Population Variability and Ecology of *Striga aspera in* Kogi State, Nigeria

**DOI:** 10.1101/2020.04.11.017574

**Authors:** Aigbokhan Emmanuel Izaka, Ohiaba Emmanuel Enemadukwu

## Abstract

Hemiparasitic *Striga* (Orobanchaceae) commonly called witchweed is native to tropical Africa. *Striga aspera* parasitizes wild grasses and its distribution range in Nigeria extends from the Sudan savanna to Guinea Savanna to the southern limit of the Derived savanna just before the forest belt is reached. This study aims to identify and delineate the incidence and distribution range and infestation patterns of *Striga aspera* within the different floristic areas within in Kogi State (Southern Guinea Savanna) and to establish if vegetation type and edaphic have potential influences on *Striga* presence. To determine the distribution range and potential hosts of *Striga aspera*, several opportunistic road reconnaissance surveys were conducted traversing six major towns in Kogi State (Kabba, Okene, Lokoja, Idah, Ayingba and Igala-mela) from July to September, 2015. Identified *Striga* infested sites were georeferenced and subjected to further vegetation analysis obtained from randomly placed 0.5 m x 0.5 m quadrats in triplicates and compared with adjoining uninfected control sites. Data for the following attributes were collected: density, relative frequency, relative density and summed dominance ratio. To isolate and determine potential Striga host, an inventory of common companion plants at infested sites were taken and screened for presence of haustorium. Edaphic soils properties were determined using standard laboratory protocols. Degree of phenotypic variability within and among the different *Striga* populations were determine using 14 morphological characters obtained from 10 randomly selected witchweed plants at each infested sites and evaluated using Principal Component Analysis and hierarchical cluster analysis. Nine sites: Idaku, Alokoina (1 and 2), Ala, Adogo (1 and 2), Ichekene, Indori and Old-Egume Road) were found to be Striga infested and all were confined to the low open woodland Southern Guinea savanna (SGS) vegetation dominated by *Daniellia/Prosopis* complex. The common *S. aspera* host was found to be *Digitaria sp.* Other companion species common at infested sites were: *Sida acuta, Centrosema pubesence, Mariscus flabelliformis, Chloris pilosa, Pennisetum pedicellatum* and *Synderella nodiflora.* Soil chemical profile reveals that *S. aspera* infestation as commonly occurs in areas acidic soils of pH ranging from 5.3 to 5.7. The Cluster analysis clearly show the similarity among *S. aspera* identified while the PCA clearly segregated the different locations where *S. aspera* was found. Findings in this study suggest that not all areas in the Derived savanna in Kogi State despite similar climatic and edaphic conditions support *Striga* infestation which showed a clustered distribution pattern. This strongly support the hypothesis that vegetation types operating at the microenvironment level may exert influences in witchweed infestation patterns.

## INTRODUCTION

*Striga* is a parasitic weed which is popularly called ‘witch weed’ attacking a wide range of crops such as cereals, legumes etc. The genus *Striga* (Orobanchacecae) contains about 41 species that are found on the African continent and parts of Asia; Africa is known to be the origin of striga. The parasite competes with the host for resources; it changes host plant architecture, and reduces the photosynthetic rate and the water-use efficiency of the host [4]

In Nigeria, three major *Striga* species have been found to be infecting crops: *S. hermonthica* (sorghum, rice, and maize), *S. aspera* (rice, wild grasses), and *S. gesnerioides* (cowpea) [6]. Other adverse effects on crops are a reduction in the ear size, plant height, stem diameter and weight of the whole plant. In addition, severe damage on roots as well as stem lodging may also be observed. Yield losses range between 10–78% especially on susceptible maize cultivars. He also reported yield losses ranging from 30 –90 % in maize, 20-100% in sorghum and millet resulting from *Striga* in Nigeria[7]. It was observed that *Striga spp* that attack cereals are distinct from those on legumes. Although 30 or more species of Striga have been described, only 5 are presently of economic importance in Africa [10]. These are, in approximate order of economical importance in Africa: *Striga hermonthica* (Del.) Benth., *Striga asiatica* (L.) Kuntze, *Striga gesnerioides* (Willd.) Vatke, *Striga aspera* (Willd.) Benth., and *Striga forbesii* (Benth). All except *S. gesnerioides* are parasites of Africa’s cereal crops such as sorghum, millet, maize, and rice. *S. gesnerioides* is a parasite on cowpea [3]. Surveys in the northern Guinea savanna of Nigeria (NGS) showed that *Striga* has remained a serious problem, attacking millet, sorghum [*Sorghum bicolor* (L.) Moench], maize [*Zea mays* (L.)] and upland rice [11]. In North-East Nigeria, over 85% of fields planted to maize and sorghum were infested with *Striga* [6].

*Striga* species exhibit variation in their mode of reproduction. *S. hermonthica, S. apara* and *S. gesneriodes* are allogamous; that is they observe cross pollination and usually rely on vectors such as bees and other agents of pollination for pollen transfer. *S. asiatica* on the other hand is autogamous; that is, it observes self pollination and so, no vectors are needed for pollination instead pollens are picked by the elongation of style and fertilization takes place [7][1][2]. *Striga aspara* is an out-crosser and frequently hybridizes with *Striga hermonthica*. It is highly likely that hybrids of these two species exist in the wild[1]. *S. aspera*, which is commonly found growing on wild grasses, has also been reported as a local pest of ‘fonio’ [*Digitaria exilis* (Kippist) Stapf.], rice and maize in the savannas of Africa. Previous Study have shown that Kogi State lies within regions in Nigeria where *Striga* infestations occur[9].

This research determines the relationship between vegetation type and prevalence of *Striga aspera* infestations, evaluate the variability in *S. aspera, S. hermonthica* and their hybrids using morphological character, specific edaphic (soil) conditions that promote the incidence of *Striga aspera* infestation, the common host of *Striga aspera* and relate these to soil fertility status.

## OBJECTIVES OF THE STUDY

The Objectives of this research are:

1. To identify the common hosts of *Striga aspera* in selected area of Kogi State
2. To determine specific edaphic (soil) conditions that promote the incidence of *Striga aspera* infestation
3. To evaluate the variability in *S. aspera, S. hermonthica* and their hybrids using morphological character
4. To show the relationship between vegetation type and prevalence of *Striga aspera* infestation in Kogi State.

## METHODOLOGY

### Area of study and study techniques

Reconnaissance field surveys were carried out between July and September, 2015 in the following six randomly selected towns and villages in Kogi State with the aid of the Map of Kogi State: Kabba, Okene, Igala-mela, Anyigba, Lokoja and Idah). Where *Striga* infestation existed, the common host and prevailing weed composition and status of the edaphic condition were evaluated. The researcher informed the farmers and the village head of his mission before entering their farmlands. Simple random sampling technique was adopted where available fields was randomly selected from the four cardinal points (north, east, west, and south) of each community. Emerged *Striga* plants was counted from each field as described by Kim [7].

### Determination of the common host of *Striga aspera*

*Striga aspera* affect most of our cereals (rice, maize, sorghum, sugar-cane) and wild grasses (*Digitaria exilis* (Kippist) Stapf.[8]. Selected cultivation areas and fields was surveyed at each identifying location. Where found, the host on which *Striga* is prevalent was determined.

### Determination of Soil Chemical Composition

In each field infected by *Striga aspera*, soil cores was taken at two depths (top soil: 0-15cm; sub soil: 15-30cm) from each quadrat and was bulked for a composite sample. The top soil samples was taken to the laboratory where they were air-dried before being subjected to physical (soil moisture) and chemical analyses (Chemical composition).

### Determination of the relationship between Vegetation type and *Striga aspera infestation*

In infected field, a ten (10) quadrats 0.5m x 0.5m was taken from plots of 100 x 100 meters in a zig-zag pattern. *Striga aspera* and plants within each quadrat was uprooted, counted and recorded. Data collected was subjected to ecological analysis to determine the density, relative frequency, relative density and summed dominance ratio according to Wirjahadja and Pancho (12). From the data, relationship between vegetation type and *Striga aspera* infestation was known.

### Evaluation of the Variability in *S. aspera, S. hermonthica* and their hybrid using Morphological characters

Quantitative data from 13 morphological character (two vegetative and 11 floral characters) was obtained at flowering from whole plants (this was possible through understanding the life cycle of *Striga* and rainfall distribution period in this location). The morphological data that was measured (in mm) were: keel diameter, leaf width, width of three fused corolla, length of: bract, calyx, calyx teeth, corolla lobe, flower, leaf, lower corolla tube, ovary, pistil and upper corolla tube[2].

### Data analysis

After the quantitative weed measurements, the density, relative density, frequency, and relative frequency, summed dominant ratio (SDR) were calculated. The data on morphological characters were analyzed using principal component analysis (PCA) and average linkage cluster analysis in order to determine the Spatial dispersion and degree of variability within locations investigated. Dendrograms was constructed based on calculated squared Euclidean distances in order to determine the level of similarities between the locations where *Striga aspera* was found. All the analysis were performed on Paleontological Statistics Software Package for education and data analysis (PAST) version 2.09.

## RESULT

### Common host of *Striga aspera*

The common host of *Striga aspera* is *Digitaria exilis*

### Determination of the relationship between Vegetation type and *Striga* infestation pattern

For Ichekene (Idah L.G.A of Kogi State): At Ichekene, *Striga aspera* was found on the road side intermingling with *Panicum maximum, Sida acuta, Ageratum conyzoides and Digitaria sp.*

**Table 1:** Quadrat count for Alokoina 1[*Striga aspera* near cassava plantation (Idah L.G.A of Kogi State)]:

| Name of Weeds | Density | Freq. | Rel. Density | Rel. Freq. | SDR |
| --- | --- | --- | --- | --- | --- |
| 1. *Digitaria sp* | 4.2 | 1 | 44.7 | 50 | 89.4 |
| 2. *Striga aspera* | 5.2 | 1 | 55.3 | 50 | 110.6 |

**Table 2:**
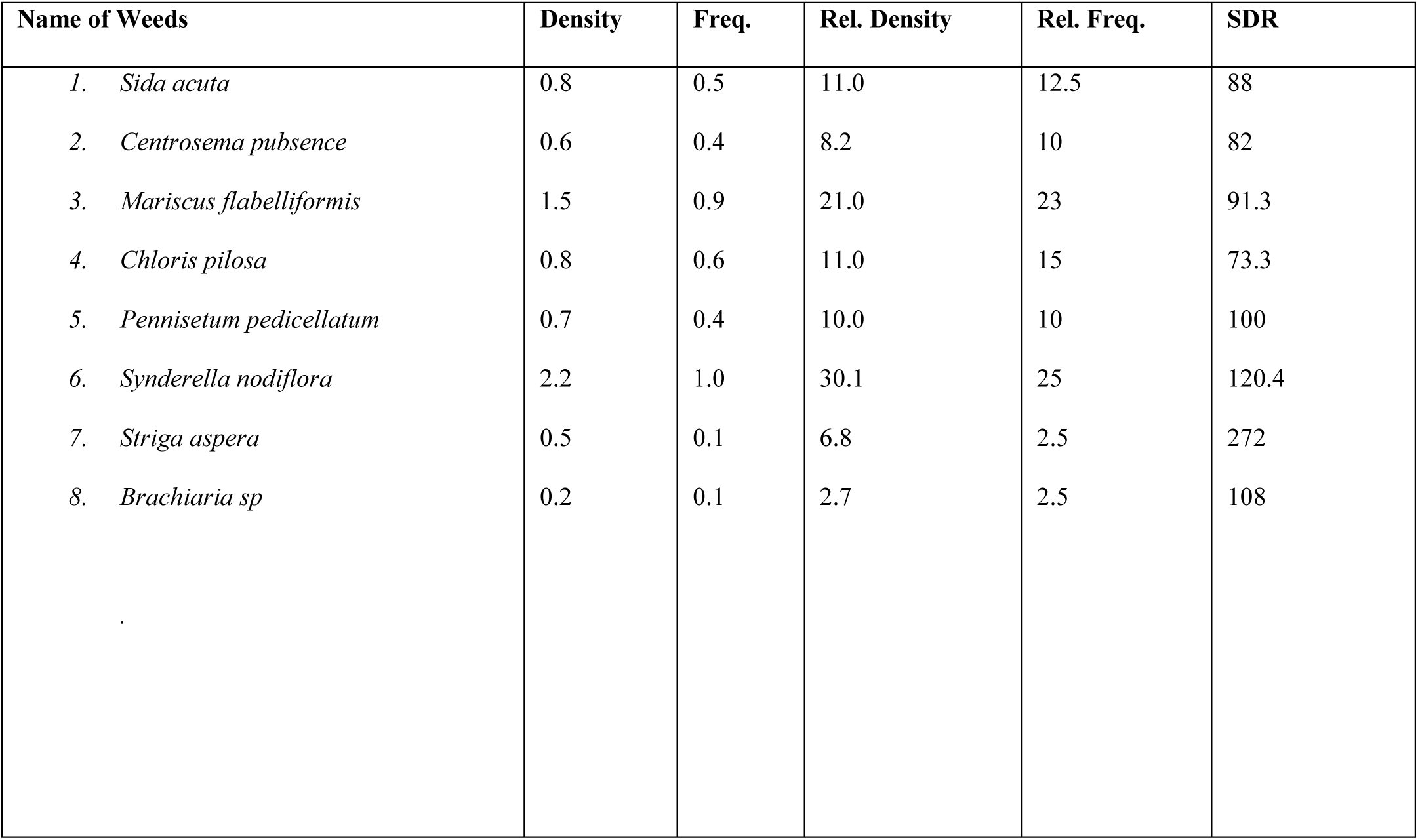
Quadrat count for Alokoina 2 (Idah L.G.A of Kogi State):

**Table 3:**
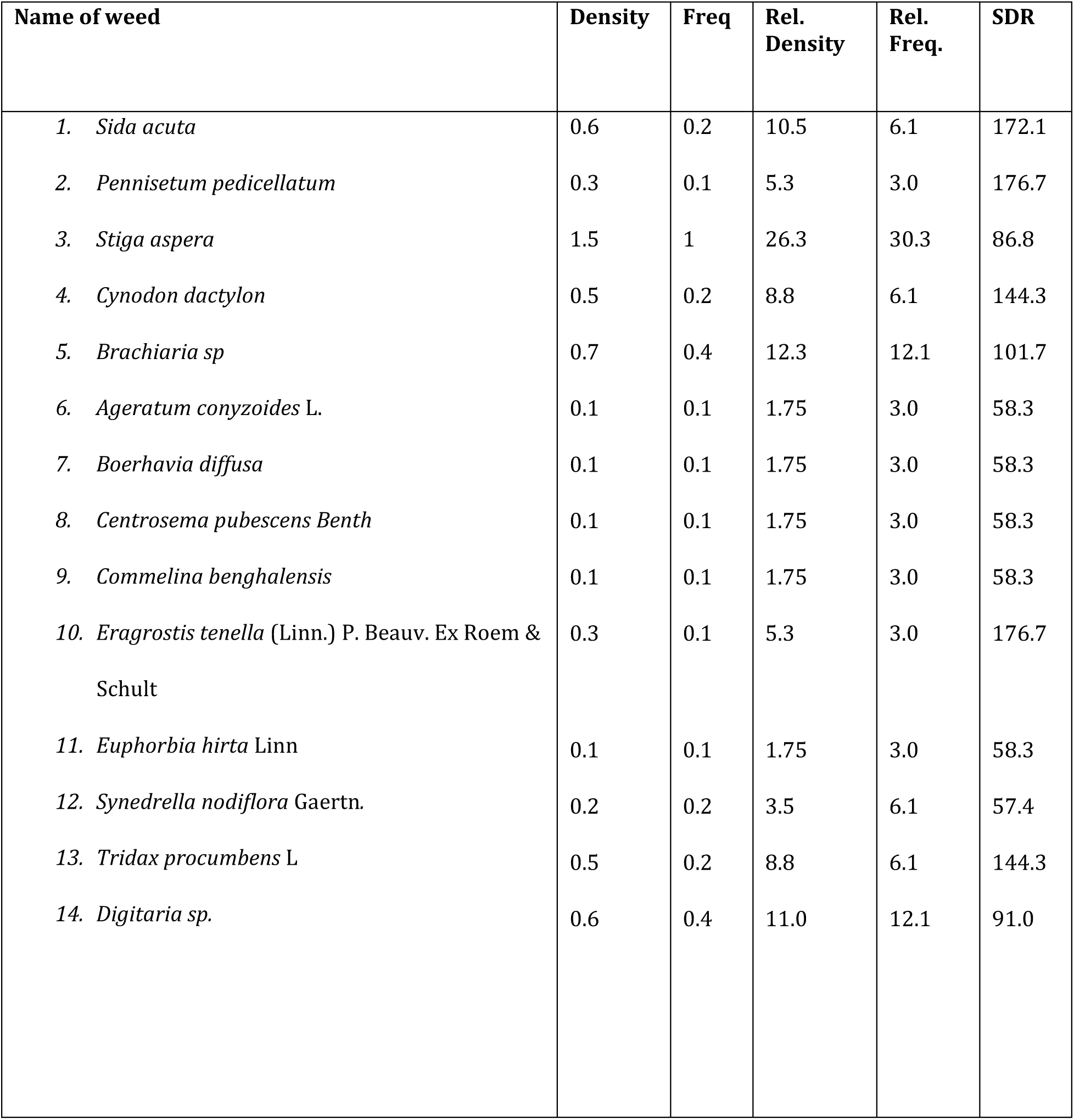
Quadrat count for Idaku (Ageokuta L.G.A of Kogi State):

**Table 4:** Quadrat count for Indori (Lokoja)

| Name of Weeds | Density | Freq. | Rel. Density | Rel. Freq. | SDR |
| --- | --- | --- | --- | --- | --- |
| 1. *Sida acuta* | 0.6 | 0.2 | 18.8 | 7.7 | 244 |
| 2. *Digitaria sp.* | 0.4 | 0.7 | 12.5 | 27 | 46.3 |
| 3. *Chloris pilosa* | 0.6 | 0.4 | 18.8 | 15.3 | 122.9 |
| 4. *Pennisetum pedicellatum* | 0.1 | 0.1 | 3.1 | 3.8 | 81.5 |
| 5. *Striga aspera* | 0.8 | 0.8 | 25 | 31 | 80.6 |
| 6. *Oldenlandia herbacea* | 0.2 | 0.1 | 6.3 | 3.8 | 165.8 |
| 7. *Synedrella nodiflora* Gaertn. | 0.2 | 0.1 | 6.3 | 3.8 | 165.8 |
| 8. *Tridax procumbens* L | 0.3 | 0.2 | 9.4 | 7.7 | 122.1 |

**Table 5:** Quadrat count for Ala (Idah L.G.A of Kogi State)

| Name of Weeds | Density | Freq. | Rel. Density | Rel. Freq. | SDR |
| --- | --- | --- | --- | --- | --- |
| 1. *Sida acuta* | 0.8 | 0.6 | 16 | 13.0 | 123.1 |
| 2. *Chloris pilosa* | 0.4 | 0.6 | 8 | 13.0 | 61.5 |
| 3. *Zea mays* | 0.6 | 0.8 | 12 | 17.4 | 69.0 |
| 4. *Striga aspera* | 1.4 | 1 | 28 | 21.7 | 129.0 |
| 5. *Spigelia anthelmia* Linn. | 0.4 | 0.5 | 8 | 11.0 | 72.7 |
| 6. *Euphorbia hirta*. | 0.6 | 0.6 | 12 | 13.0 | 92.3 |
| 7. *Tridax procumbens* L | 0.8 | 0.5 | 16 | 11.0 | 145.5 |

\* Rel. Density-Relative Density, Rel. Freq-Relative Frequency, SDR-Summed Dominant Ratio

For Old Egume, a total of 9 *Striga aspera* was found intermingling with *Centrosema pubsence, Imperata cylindrical, Cynodon Datcylon, digitaria sp.*

For Adogo 1 and 2, *Striga aspera* was found on the road side intermingling with *Cynodon dactylon, Sida acuta, Digitaria sp., Panicum maximum*

### Result of Physiochemical analysis of soil samples gotten from the field

**Table 6:**
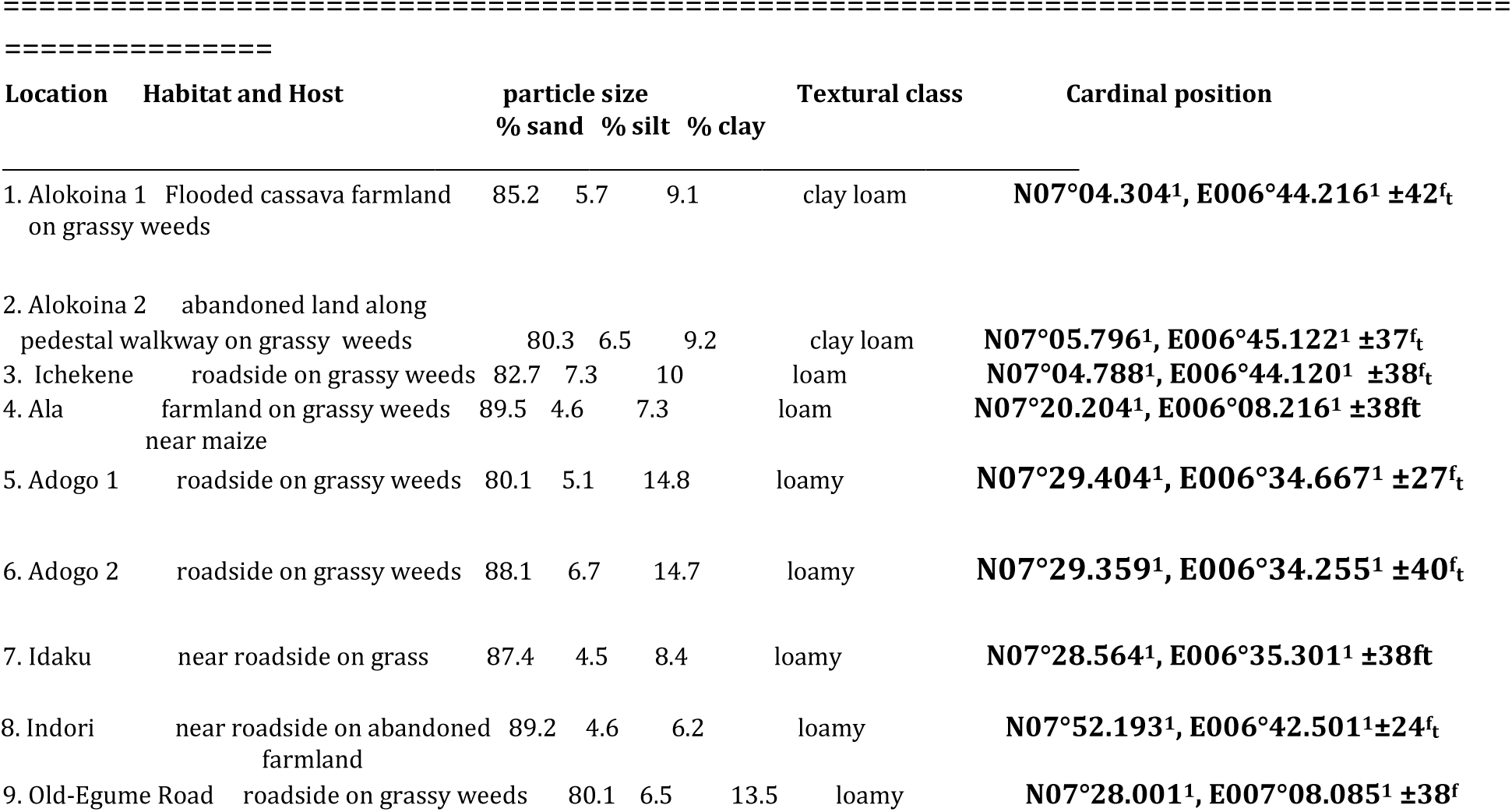
Soil physical properties, host and habitat of 9 *Striga aspera* collection sites in Kogi State.

**Table 7:**
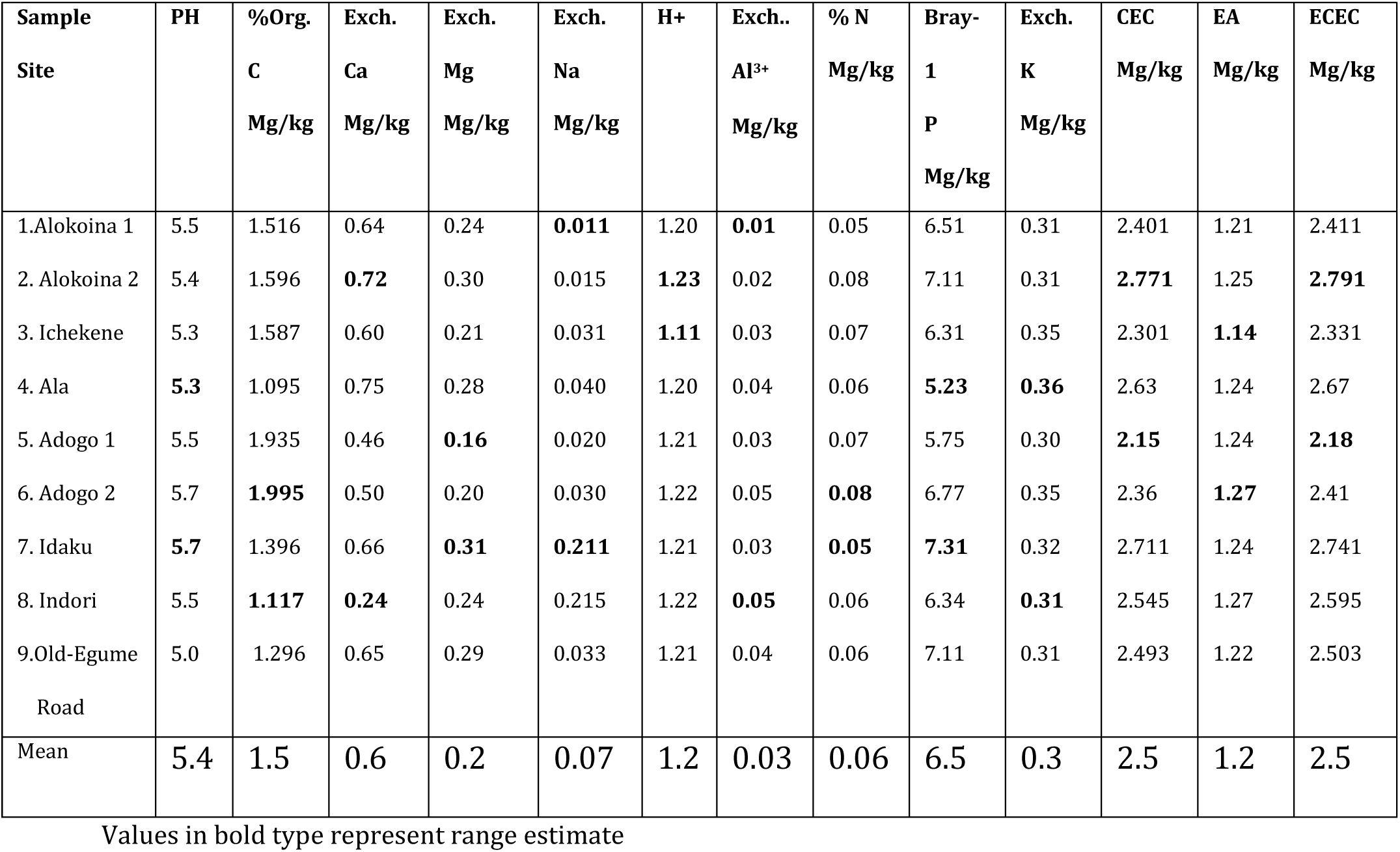
Soil chemical properties of 9 *Striga aspera* collection sites in Kogi State, Nigeria.

The result chemical analysis shows that the PH of the soil to be acidic with mean PH of 5.4 and with range (5.3-5.7); mean of %Organic matter is 1.5 (range 1.1-2.0); mean of Exchangeable Calcium is 0.6 with range (0.2-0.7); mean of Exchangeable Magnesium is 0.2 with range (0.2-0.3), mean of Exchangeable Sodium is 0.07 with range (0.01-0.21); mean of H^+^ is 1.2 with range (1.1-1.2); mean of Exchangeable Aluminum is 0.03 with range (0.1-0.5); mean of %Nitrogen is 0.06 with range (0.50.8); mean of Bray-1Phosphorus is 6.5 with range (5.2-7.3); mean of Potassium is 0.3 with range (0.3-0.4); mean of Cation Exchangeable Capacity is 2.5 with range (2.2-2.8); mean of Exchangeable Acidity is 1.2 with range (1.1-1.3) and finally, mean of Effective Cation Exchangeable Capacity is 2.5 with range (2.2-2.8).

### Result of the measurement of Morphological characters (mm) of *Striga aspera* analyzed statistically using Principal Component Analysis

#### Spatial Dispersion

Scatter plots from morphological characteristic of *S. aspera* for each locations onto the principal components is shown in Figures 1. The dispersion pattern clearly segregated the different locations where *S. aspera* was found. The PCA loadings, indicate that dispersion along PC1 axis was mainly influenced mostly due to the presence of the locations: Adogo, Old Egume, Alokoina, Idaku, Indori or absence of the locations: Ala, Ichekene while the presence of Locations: Indori, Ala, Alokoina, Ichekene or absence of locations: Old-Egume, Idaku, Adogo accounted for dispersion observed along PC2 axis.

**Figure 1:**
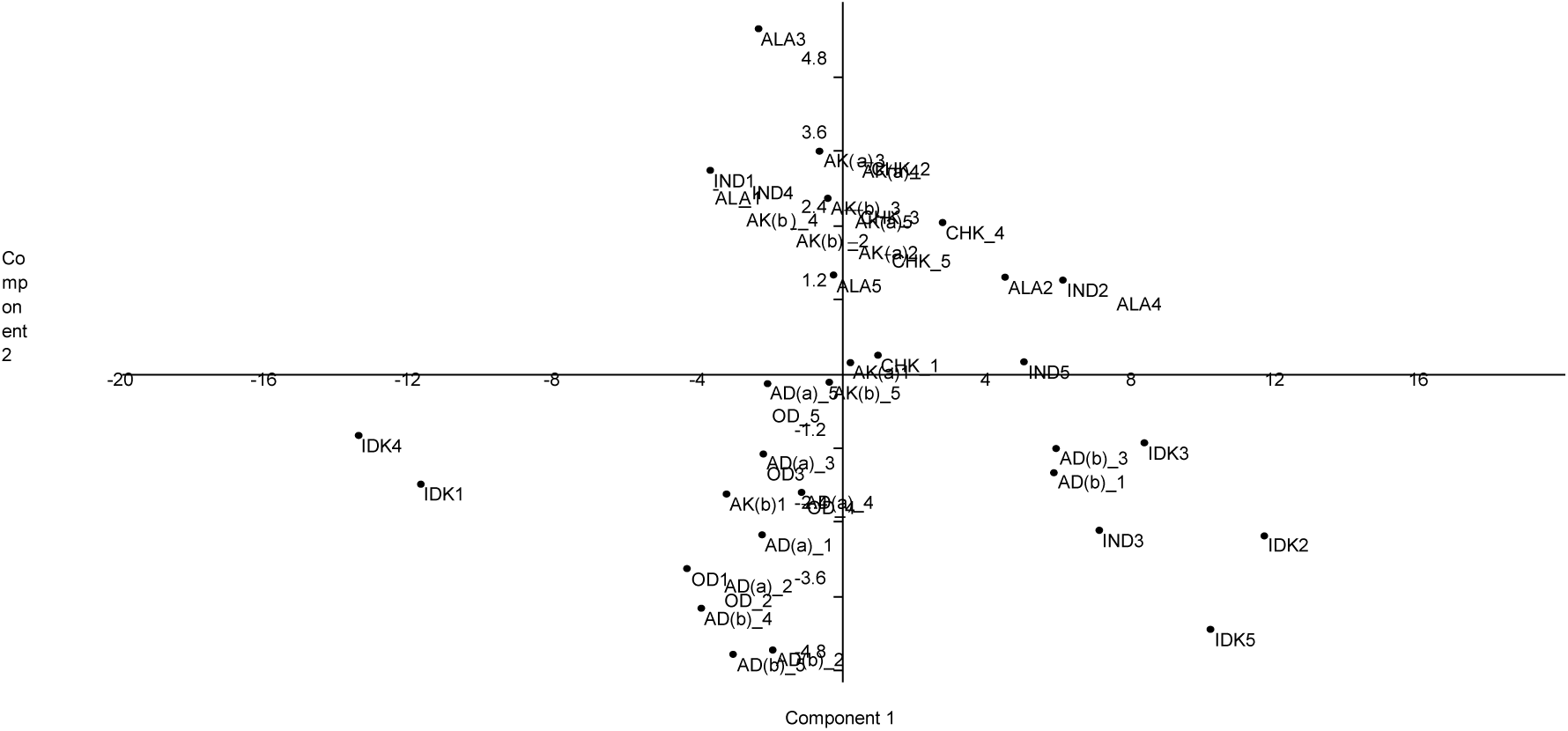
Principal components analysis comparing spatial displacement of the different locations where *S. aspera* was found based on the morphological characteristics of *S. aspera* measured for each location. Where CHK= Ichekene; AK(a)= Alokoina Site 1; AK(b): Alokoina Site 2; IND=Indori; IDK=Idaku; AD(a)=Adogo site 1, AD(b)=Adogo site 2, OD=Old Egume Rd and ALA= Ala

#### Spatial Dispersion

Spatial Dispersion of the locations identified in this studies and comparing it with the work of Aigbokhan *et. al.* [2]. The dispersion in Fig. 2 shows that the *Striga* from the Northern region [2] is morphologically different from the one identified in this work but I was unable to compare if there is any variability in *S. aspera /S. hermonthica* because *S. hermonthica* was not encountered in the survey.

**Figure 2:**
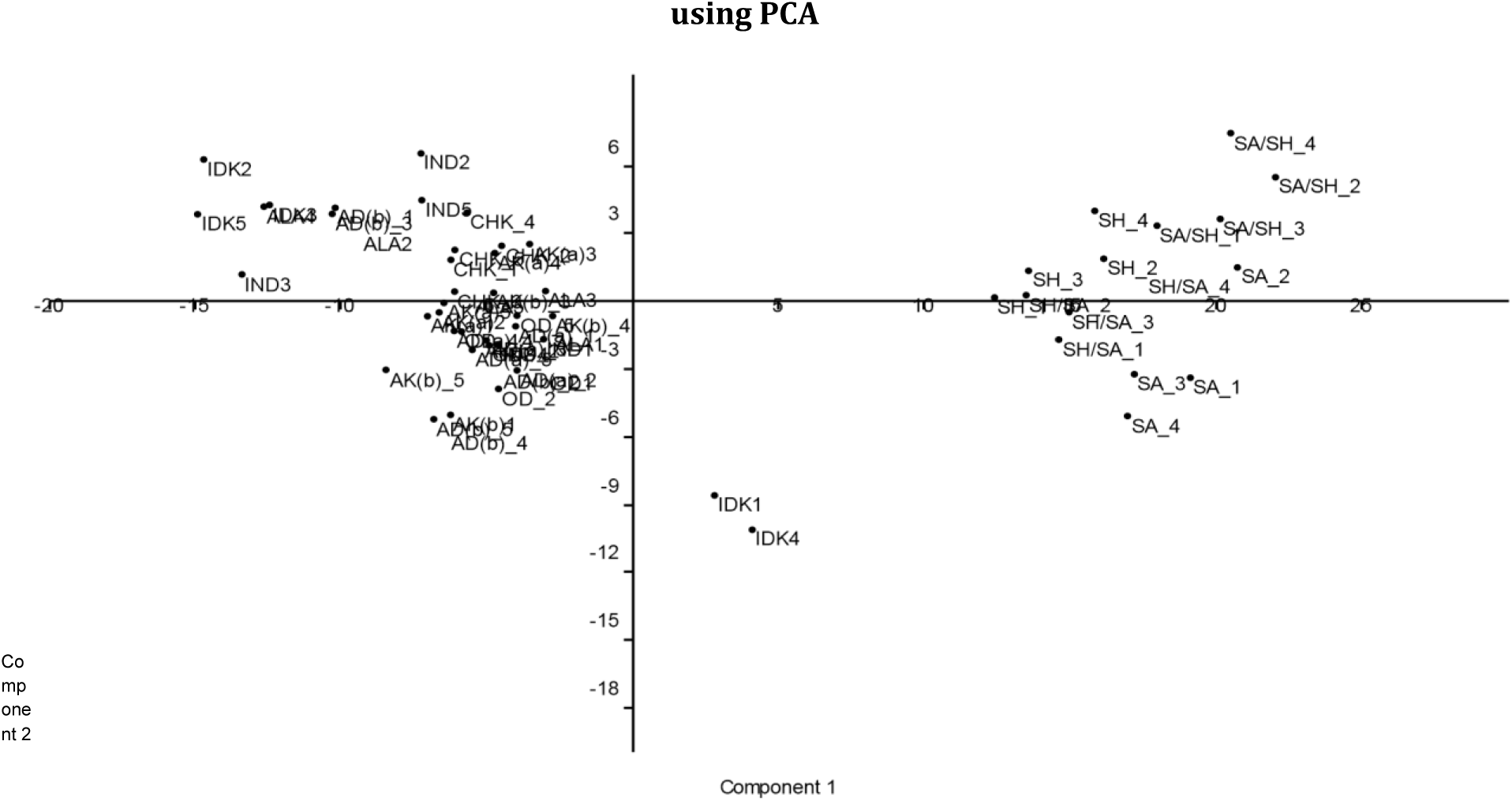
Principal components analysis comparing spatial displacement of the different locations where *S. aspera* identified in this work and the comparison with the work of Aigbokhna *et. al.* (2). Where SH= *Striga hermonthica*, SA= *Striga aspera*

### Comparing the Morphological data with the work of Aigbokhan *et. al.* [2) and checking for the variability

#### Similarity Profile

The similarity profile matrix generated using the Bray-Curtis method for 9 locations shows the Hierarchical cluster in Figure 3. This reveal that some morphological data for some sites shows close uniformity and some morphological data for some sites does not show any close uniformity. Both the AD(b) 4, 5 were most similar at nearly 0.94 similarity index level with Idaku being most unique and markedly different from all the other locations which shared similarity with other locations only at nearly 0.84 similarity index.

**Figure 3:**
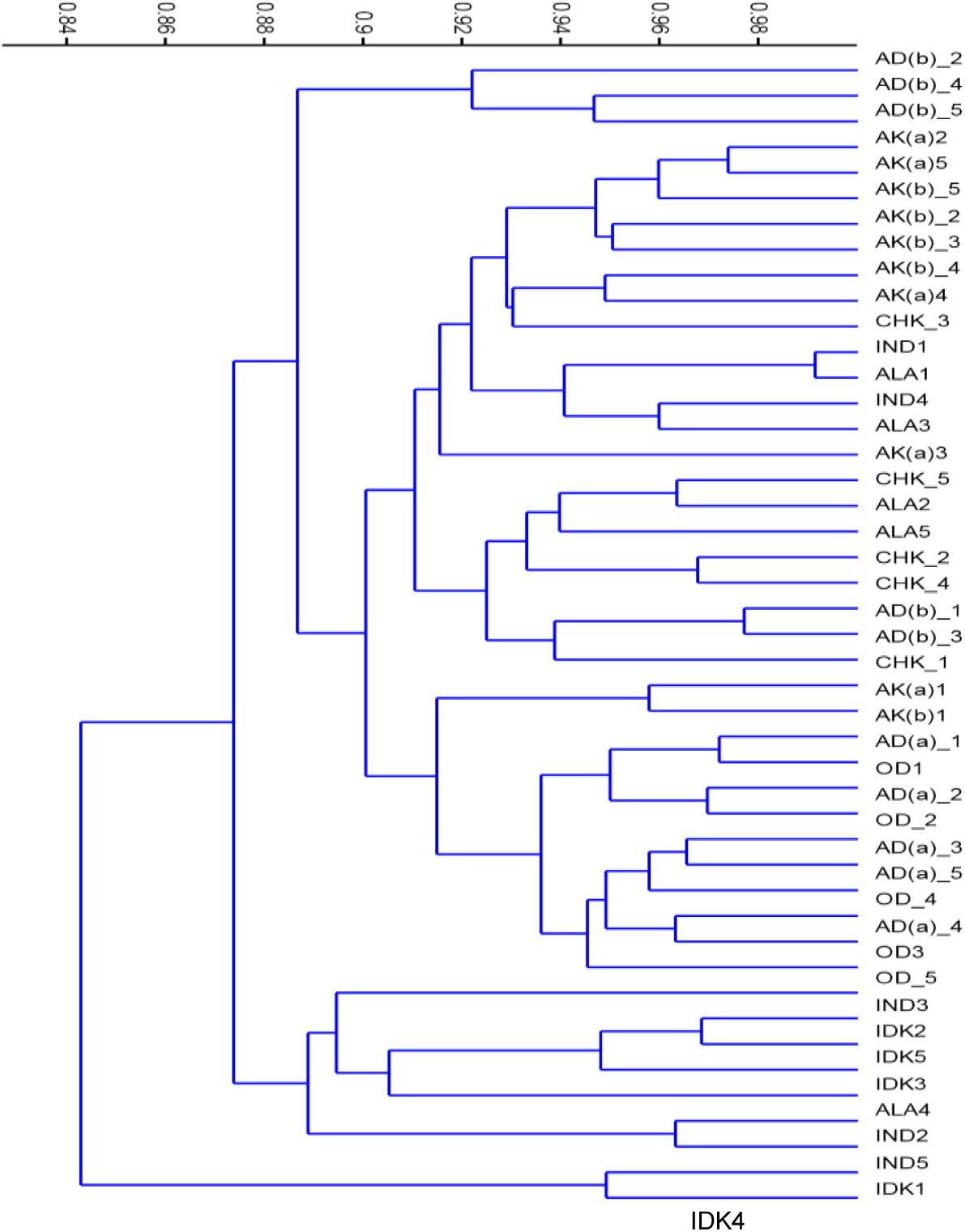
A cluster diagram showing position of different site of *S. aspera* based on similarity estimates based on Bray-Curtis index using paired group algorithm. The cut off point is set at 0.82

#### Similarity Profile

The similarity profile matrix generated using the Bray-Curtis method for 9 locations and with the work of Aigbokhan *et. al.* (2) shows the Hierarchical cluster in Figure 4. This reveal that there is no close uniformity between the morphological data for these work and the work of Aigbokhan *et. al.*, (2) but they only shared similarity at 0.73 similarity index.

**Figure 4:**
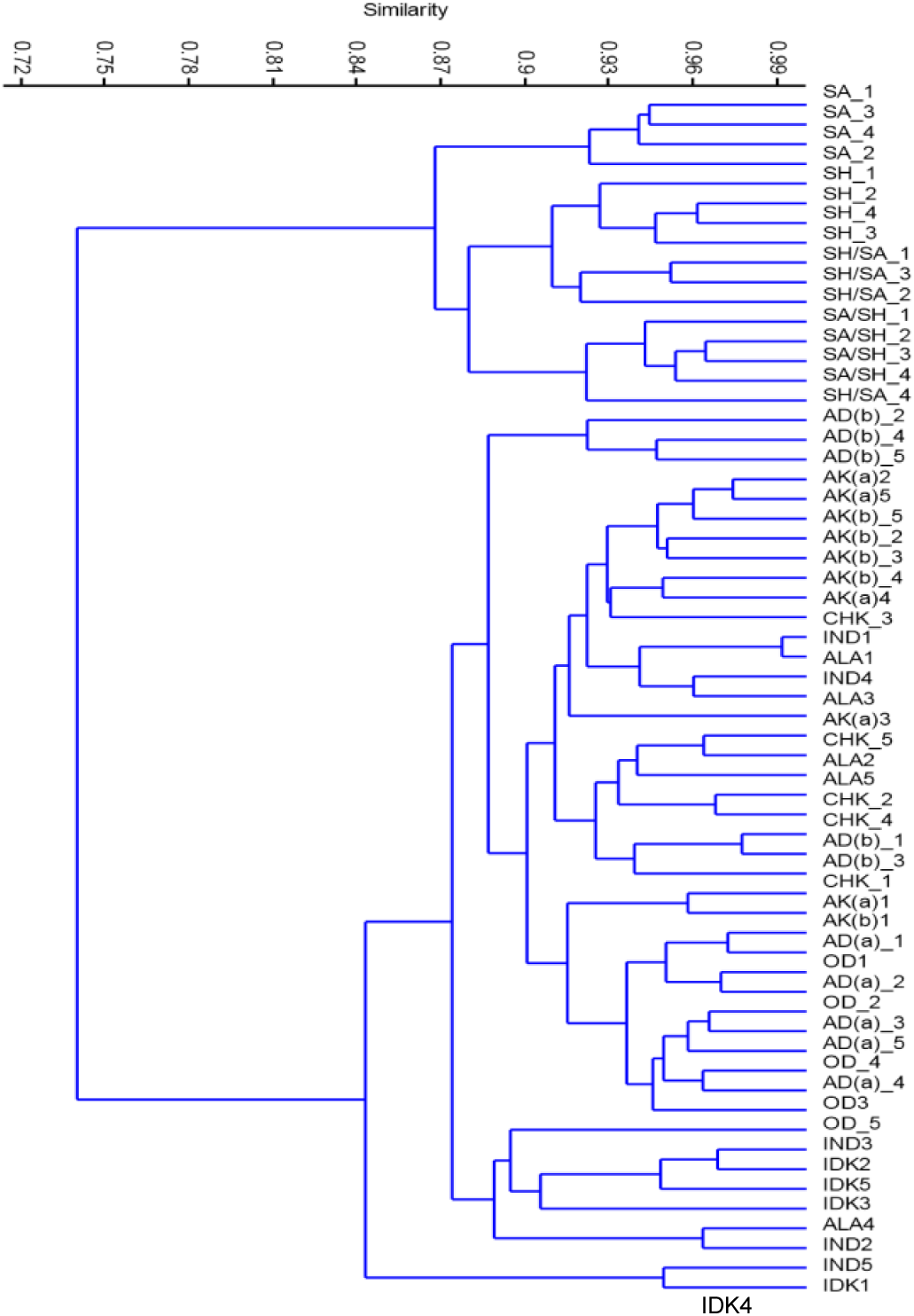
A cluster diagram (Dendrogram) showing the comparison between different sites identified for *S. aspera* in the survey and the work of Aigbokhan *et. al.* (2) based on similarity estimates based on BrayCurtis index using paired group algorithm. The cut off point is set at 0.72

**Figure 5:**
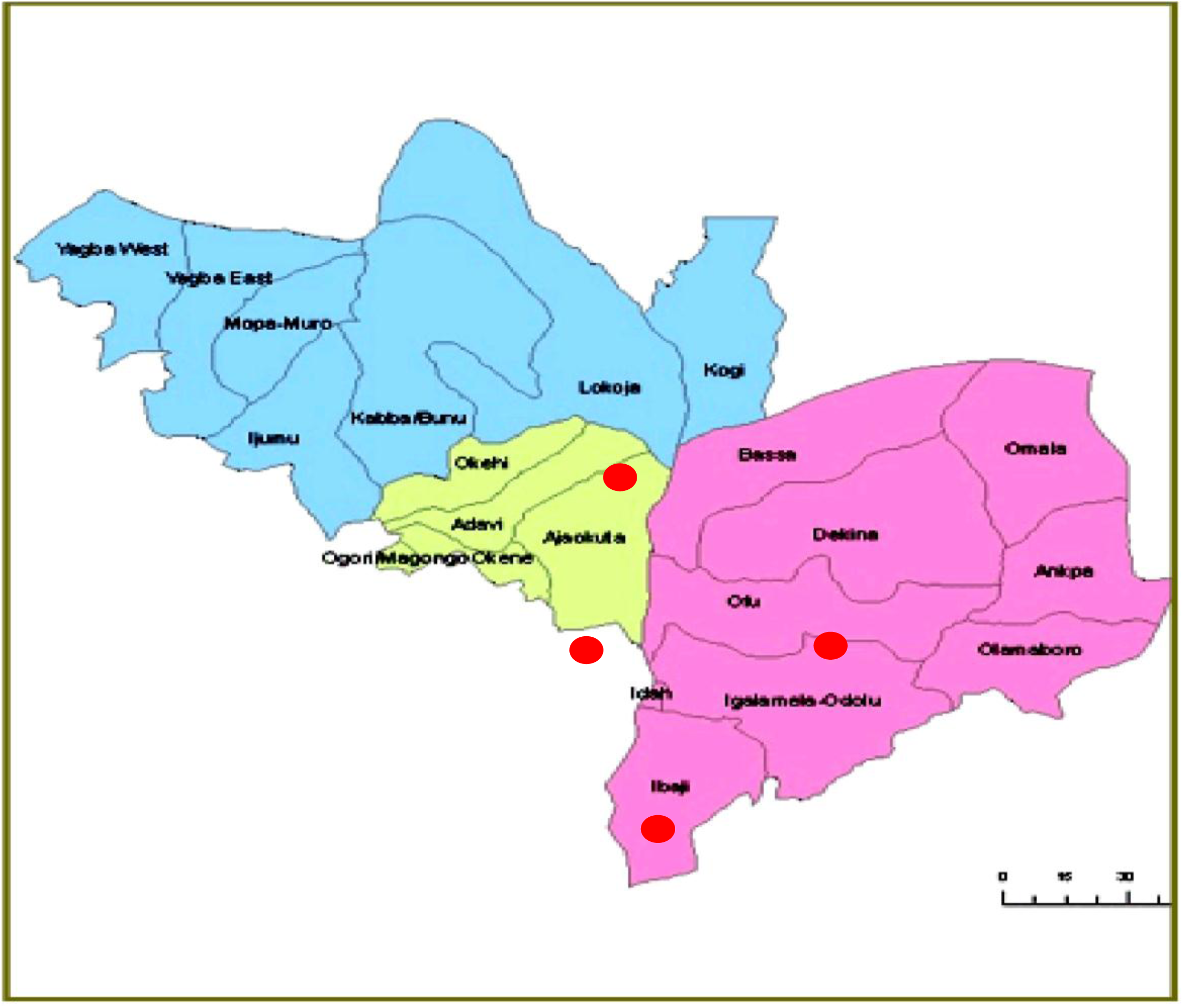
MAP OF KOGI STATE SHOWING THE LOCAL GOVERNMENT AREAS (AREAS WITH SHOWS 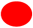 AREAS WITH *STRIGA ASPERA* INFESTATION)

**Figure 6:**
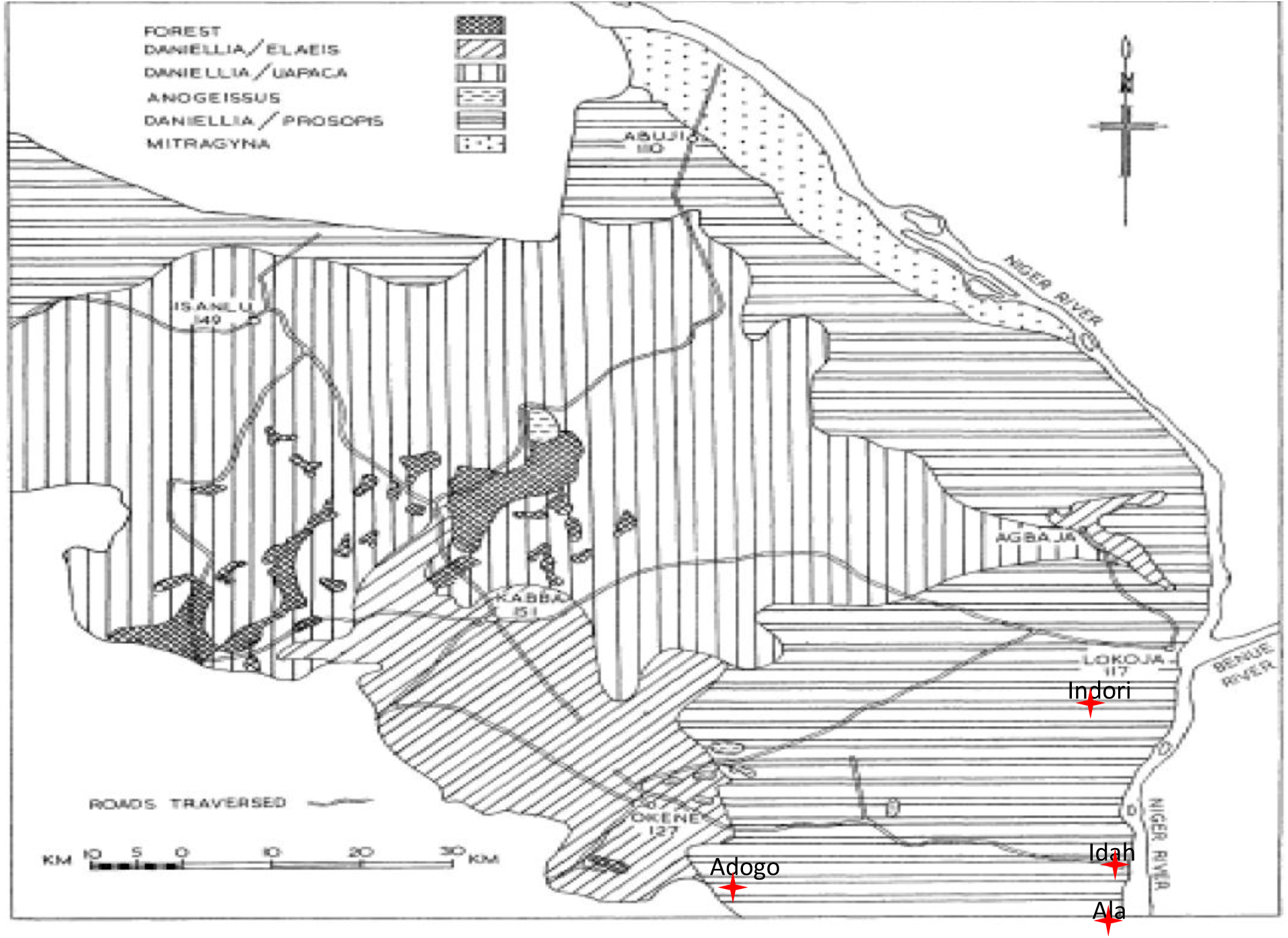
Vegetation Map of Kabba. Places of *Striga aspera* infestation is represented on the map with 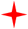

## DISCUSSION AND CONCLUSION

The incident of first seedling emergence and some structural characteristics of the seedlings are unique to the life cycle of *S. aspera and S. hermonthica.* Ramaiah called the trilobed plumule on the elongated neck in *S. hermonthica* as “*Striga* initial”[10]. This was the first report to distinguish between *S. aspera and S. hermonthica* seedlings by the presence of hairs on *S. aspera* shoot premordia. Though, *S. hermonthica* was not found in the survey but the uniqueness of their character being pointed out by Ramaiah *et. al.* (10) has given me an ample opportunity to identify *S. aspera* in the field.

Mohammed identified leaf size, flower density, inflorescence length, habitat type, presence and absence of flower stalk, equality of calyx teeth and number of calyx ribs as the major characters separating *S. aspera and S. hermonthica* [8].. Cluster analysis between this work and the work of Aigbokhan *et. al.* (2) based on morphological characters shows a relatively low 0.72 level of similarity between the two work.

The common host of *Striga aspera* is *Digitaria exilis.* In the field, *S. apera* was found intermingling with different species like *Sida acuta, Centrosema pubsence, Mariscus flabelliformis, Chloris pilosa, Pennisetum pedicellatum, Synderella nodiflora* etc apart from its host *Digitaria sp.*

The distribution of *Striga aspera* in selected locations in Kogi state has been accessed. Although, it is highly likely that hybrids of these two species exist in the wild (1), the present study only spotted *S. aspera.*

The result from chemical analysis of the soil reveals that *S. aspera* was predominantly found in acidic soil with mean PH range from 5.3-5.7, indicating an acidic soil condition. Also mean of % Organic matter is 1.5 (range 1.1-2.0); mean of Exchangeable Calcium is 0.6 with range (0.2-0.7); mean of Exchangeable Magnesium is 0.2 with range (0.2-0.3), mean of Exchangeable Sodium is 0.07 with range (0.01-0.21); mean of H^+^ is 1.2 with range (1.1-1.2); mean of Exchangeable Aluminum is 0.03 with range (0.1-0.5); mean of % Nitrogen is 0.06 with range (0.5-0.8); mean of Bray-1Phosphorus is 6.5 with range (5.2-7.3); mean of Potassium is 0.3 with range (0.3-0.4); mean of Cation Exchangeable Capacity is 2.5 with range (2.2-2.8); mean of Exchangeable Acidity is 1.2 with range (1.1-1.3) and finally, mean of Effective Cation Exchangeable Capacity is 2.5 with range (2.2-2.8). These factors are important in monitoring the species distribution.

Morphological characters are very commonly used to assess relationships between species. Scatter plots from morphological characteristic of *S. aspera* for each locations onto the principal components is shown in Figures 1. The dispersion pattern clearly segregated the different locations where *S. aspera* was found. The PCA loadings, indicate that dispersion along PC1 axis was mainly influenced mostly due to the presence of the locations: Adogo, Old Egume, Alokoina, Idaku, Indori or absence of the locations: Ala, Ichekene while the presence of Locations: Indori, Ala, Alokoina, Ichekene or absence of locations: Old-Egume, Idaku, Adogo accounted for dispersion observed along PC2 axis.

The distribution of *Striga aspera* has been identified. The common host of *Striga aspera* is *Digitaria exilis.* In the field, *S. apera* was found intermingling with different species like *Sida acuta, Centrosema pubsence, Mariscus flabelliformis, Chloris pilosa, Pennisetum pedicellatum, Synderella nodiflora* etc apart from its host *Digitaria sp.* Comparative morphological analysis which was done using Principal Component Analysis (PCA) / Cluster analysis shows variability between the findings of Aigbokhan *et. al.* (2) on morphological characteristics and the morphological characteristics measured in this work. Findings in this study suggest that not all areas in the Derived savanna in Kogi State despite similar climatic and edaphic conditions support *Striga* infestation which showed a clustered distribution pattern. This strongly support the hypothesis that vegetation types operating at the microenvironment level may exert influences in witchweed infestation patterns.

